# Impacts and mechanisms of CO_2_ narcosis in bumble bees: Narcosis depends on dose, caste and mating status and is not induced by anoxia

**DOI:** 10.1101/2022.07.01.498474

**Authors:** Anna Cressman, Etya Amsalem

**Affiliations:** Department of Entomology, Center for Chemical Ecology, Center for Pollination Research, Huck Institutes of the Life Sciences, Pennsylvania State University, University Park, PA 16802, USA

**Keywords:** carbon dioxide, insects, reproduction, metabolism, anesthesia

## Abstract

Carbon dioxide (CO_2_) is commonly used to immobilize insects and to induce reproduction in bees. However, despite its wide use and potential off-target impacts, its underlying mechanisms are not fully understood. Here we used *Bombus impatiens* to examine whether CO_2_ impacts are mediated by anoxia and whether these mechanisms differ between female castes or following mating.

We examined the behavior, physiology, and gene expression of workers, mated and virgin queens following exposure to anoxia, hypoxia, full and partial hypercapnia, and control. Hypercapnia and anoxia caused immobilization, but only hypercapnia resulted in behavioral, physiological, and molecular impacts in bees. Recovery from hypercapnia resulted in increased abdominal contractions and took longer in queens. Additionally, hypercapnia activated queens’-but inhibited workers’ ovaries in a dose-dependent manner and caused a depletion of fat-body lipids in both. All responses of hypercapnia were weaker following mating in queens. Analysis of gene expression related to hypoxia and hypercapnia supported the physiological findings in queens, demonstrating that the overall impacts of CO_2_, excluding virgin queen ovaries, were unique and were not induced by anoxia. This study contributes to our understanding of the impacts and the mechanistic basis of CO_2_ narcosis in insects and its impacts on bees’ physiology.

## Introduction

Carbon dioxide (CO_2_) is a prevalent gas present in the atmosphere that can vary in concentration in different microclimates. Low concentrations of CO_2_ are often used as an attractant or repellent and are identified by designated receptors in insects (1). They can guide host plant finding and increase foraging efficiency when flowers are open in herbivores (2), nest digging in ants (3, 4) and nest maintenance behaviors in social bees and termites (5, 6). In contrast, hypercapnic concentrations of CO_2_ can result in pervasive physiological, molecular, and behavioral changes (2, 7). Full hypercapnia (100% CO_2_) is a commonly used anesthetic during experimental procedures in insects but may result in unwanted side effects. Despite its wide use and importance in insects, the mechanisms underlying CO_2_’s mode of action are not fully understood.

CO_2_ narcosis has pleiotropic impacts on insect physiology and behavior. These effects have been documented in multiple orders including Orthoptera (8), Diptera (9, 10), Hemiptera (11), and Hymenoptera (12–15). CO_2_ narcosis in *Drosophila melanogaster* resulted in changes in fecundity and longevity (10) and reduced climbing behavior (16). CO_2_ also inhibited feeding and drinking in *Empoasca devastans* (11), and reduced social aggregation in *Locusta migratoria* (8, 16). In addition, CO_2_ treatment has specific impacts on insect reproduction, some are being used for practical purposes, such as initiating egg laying in queen bees (17, 18). However, these impacts are not limited to stimulation and may also include suppression of reproduction in some species. For example, exposure to CO_2_ led to a reduction in insemination frequency in *Glossina* females (19), suppressed ovarian development in adult *Tribolium castaneum* (20), reduced egg laying in *Pyrrhocoris apterus* (21), and inhibited ovary activation in honey bee workers (14, 15, 22). Similar changes following CO_2_ were found in the gene expression profile of multiple species, demonstrating changes in genes related to vitellogenesis (23–25), insulin and juvenile hormone (JH) signaling pathways (23) and oxidative stress (14, 23).

Earlier studies examining the mechanisms underlying CO_2_ suggested that it affects the nervous system through rapid changes in intracellular pH (2). While the membrane acidity was shown to increase following CO_2_ in numerous species (26–28), it has never been directly linked to the unique effect of CO_2_ and can be caused by several potential mechanisms. One of these is hypoxia, whereby CO_2_ operates via the lack of oxygen (2, 29–31). This too could lead to acidic intracellular pH; however, the data are controversial. In termites, both CO_2_ and hypoxic conditions increased the reproductive output in *Reticulitermes speratus* (32) and also induced similar changed in reproductive-related genes (33). On the other hand, a nitrogen treatment (anoxia) failed to induce a shift from a gregarious to a solitary state in *Locusta migratoria*, as did CO_2_ (8). The same was found in crayfish where a nitrogen treatment did not produce the same behavioral (avoidance response) and physiological (increase in heart and ventilatory rates) responses caused by CO_2_ (34). Finally, in cricket larvae, some of the impacts of CO_2_ could be mimicked by nitrogen (feeding and drinking inhibition), but not all of them (e.g., metabolic rate) (29).

Another hypothesis proposed that CO_2_ operates directly on the central nervous system, but how exactly this works and whether the mechanism is conserved across species remain open. Studies in *Drosphila melanogaster* and crayfish demonstrated that CO_2_ blocks glutamate receptors at the neuromuscular junction (NMJ) and through inhibition of recruiting motor neurons within the CNS (28, 35). In crayfish, NMJ responds to exogenous glutamate and to saline that was adjusted to pH 5.0 by depolarization. However, the synaptic transmission is shut down in response to CO_2_-saturated water and does not respond to either exogenous glutamate or to acidic pH (5.0). Similar evidence for CO_2_ blocking the glutamate receptors were obtained also in Drosophila (28). There, it was also shown that CO_2_ has actions not only in the CNS, but also in the periphery.

Additional studies provided more information about the cascade process between CO_2_ administration and its downstream effects. First, CO_2_ narcosis is followed by changes in brain biogenic amines. Levels of dopamine, tryptophan, and tyrosine in pollen-fed honey bee workers were significantly reduced following CO_2_ compared to untreated controls, whereas in caged queens, dopamine levels decreased in 45% (36). Levels of octopamine, content of cAMP, and sensitivity of their receptors were modified also in *Locusta migratoria* following CO_2_ narcosis (30). Second, JH, a gonadotropin in most insects, was shown to spike following CO_2_ narcosis in bumble bees (23), and when young queens were fed with JH inhibitor, CO_2_ had no effect on their ovaries and fat body macronutrient amounts as compared to controls (Barie, Levin, Amsalem, submitted). Combined, these changes likely result from the neural changes and the decrease in intracellular pH caused by CO_2_ and lead to the diverse behavioral and physiological effects observed downstream. However, the cascade of reactions and the overall mechanism are not fully understood.

In this study we examined whether CO_2_ causes immobilization and transition to reproduction by inducing anoxia and whether these mechanisms differ between female castes or following mating using the social bumble bee *Bombus impatiens*. This bee goes through an annual life cycle where a single mated queen produces workers until the nest reaches several hundreds of bees, and then switches to produce virgin queens that will mate and go into a winter diapause (37). The effects of CO_2_ were extensively studied in bees where it induced a transition to reproduction in honey bee and bumble bee queens (17, 18) and inhibition of reproduction in honey bee workers (38). It was further shown to induce a metabolic shift in bumble bee queens, reducing lipid levels in the fat body and increasing glycogen and protein in the ovaries (14, 23) (Barie, Levin, Amsalem, submitted), as well as increasing the hemolymph levels of JH (23). The effects of CO_2_ on queen metabolism persist even when queen’s ovaries were removed but did not persist when queens were fed with a JH inhibitor (Barie, Levin, Amsalem, submitted), suggesting that CO_2_ impacts are mediated via JH and its effect on reproduction is secondary. It was also shown that when combined with cold storage, the CO_2_’s effect on reproduction in queens diminished with the length of cold storage (39), showing once again that the effect on reproduction is a byproduct of other processes. CO_2_ also increased aggression and flight behavior in virgin queens (14) and changed the gene expression profile of the queen’s following administration (14, 23). However, data on CO_2_ impacts throughout the different queen life stages and in workers are lacking. Altogether, bumble bees are an excellent system model to examine the mechanisms underlying CO_2_’s mode of action.

To examine whether CO_2_ is induced by anoxia, we assigned workers, virgin, and mated queens to anoxia, hypoxia, and full and partial hypercapnia, as compared to controls and examined the behavior, physiology, and gene expression of bees. We hypothesize that if full hypercapnia and the lack of oxygen (anoxia) have different effects on bees, then CO_2_ is acting uniquely. However, if hypercapnia and anoxia have the same effects, then CO_2_ is operating by inducing anoxia.

## Methods

### Bees and treatments

Colonies were obtained from Koppert Biological Systems (Howell, MI) and maintained in environmental chambers under darkness, 28-30° C, 60% relative humidity, and supplied with *ad libitum* 60% sugar solution and fresh pollen purchased from Koppert. Newly-emerged workers and gynes (< 24 hours old and distinguished by their silvery appearance) were collected from multiple source colonies (n=12), tagged with a colored number, and randomly assigned a cage (11 cm diameter and 7 cm high). Workers were kept in groups of three while gynes were kept alone. Cages were randomly assigned to one of five gas treatments including full hypercapnia (FH, 100% CO_2_), partial hypercapnia (PH, 50% CO_2_, 50% N_2_), hypoxia (HY, 14% O_2_, 86% N_2_), anoxia (AN, 100% N_2_), and control (C, no gas exposure). Gas treatments were conducted for one full minute of air flow into a nearly sealed plastic cage. Queens and workers typically lose mobilization within seconds when exposed to full hypercapnia, partial hypercapnia, and anoxia, but do not lose mobility in hypoxic or control conditions. The seal was left on the container for an additional 30 minutes after exposure, allowing the cage to reach an equilibrium with the ambient air and was removed afterwards. Following treatments, cages were placed back in the environmental chamber.

#### Workers

534 newly-emerged workers (the split of sample size to treatments and variables is provided in Table S1) were collected and kept in group of three. In the first experiment (“single treatment”), workers (n=309) were treated with gas on day one of age and were kept for six- and ten-days post treatment. Following recovery from the gas treatment, workers were immediately observed for abdominal contractions and for aggressive behavior between nestmates for 20 min per day on days 1-6. They were then stored in −80 °C until further analyses. Six-day-old workers were examined for ovary activation, fat body lipid amount, and head gene expression, and 10-day-old workers were examined for total number of eggs laid per cage. The second experiment with workers (“multiple treatments”, n=225) was identical to the first one with two differences: first, workers were treated daily on days 1-6 of age to account for age of exposure being a possible factor in the effectiveness of the gas treatment, and second, workers were frozen at the age of 7 instead of 6 days to allow them more time to activate their ovaries. The workers of the second experiment were examined for ovarian activation, fat body lipid amount and head gene expression.

#### Queens

Newly emerged queens (n=192, Table S1) were kept in individual cages for six days and were introduced into a mating arena at the age of 6-11 days. Queens were put back in their individual cages following mating (mated) or unsuccessful attempt to mate (virgin). Mating procedure is described below. Queens were randomly assigned to a gas treatment 24 hours post-mating or the attempt to mate and were observed for abdominal contractions immediately after recovery. Queens were kept for additional ten days post treatment, resulting in all queens being 17-22 days old at the completion of the experiment. Queens were stored in −80 °C and were examined for ovary activation, egg laying, fat body lipids and head gene expression.

### Mating

At the age of six days, queens were placed in a mating arena (35 × 12 × 12 cm) with 3-4 males for every queen. Queens were given a three-hour window to mate, during which they were constantly observed. If queens did not mate, a similar attempt was conducted daily until a successful mating was observed or the queen reached the age of 11 days. This range was chosen based on the optimal age for mating found in our previous study (40). The arena was checked every 15 minutes to observe for mating pairs and pairs were removed while mating and placed back into an individual cage. The pair remains connected in their genitals for approximately 30 minutes and males were removed from the cage upon the completion of mating. Whenever a mated queen was sampled, a same-age unmated queen (virgin) was removed from the arena and sampled as well.

### Abdominal contractions

Abdominal contractions (AC) are performed by insects to facilitate gas exchange throughout the body (41, 42), and typically start upon recovery from anesthesia. The number of abdominal contractions was video-recorded and quantified for 20 minutes in queens and workers from the first movement and only in treatments that caused immobility (full hypercapnia, partial hypercapnia, and anoxia). We also quantified the time it took bees to enter anesthesia (in seconds) and the recovery time from anesthesia (in minutes). Recovery time differed between treatments and between queens and workers (see results), and therefore recording began at different timepoints after the gas treatments.

### Aggression in worker groups

Aggressive behavior in workers was observed on days 1-6 following exposure in workers of experiment 1 (single treatment). Each cage was observed for 20 minutes per day under red light between 8:00-12:00 AM. We recorded and summed four aggressive behaviors typically seen in queen-less groups of *Bombus impatiens* (14): humming (a series of wing vibrations lasting less than 3 s), climbing (one bee crawls on top of another bee), darting (a sudden movement in the direction of another bee) and attacking (biting, pushing, dragging by legs or wings, and an attempt to sting). The identity of the bee performing and receiving each behavior was recorded. The total aggressive behaviors performed by individual workers during days 1-6 was used in further analyses. Queens were not observed for aggression since they were kept alone.

### Egg laying

Cages of workers at the age 10 days (first experiment) and queen cages were checked daily for newly constructed egg cells. Ten days after the gas treatment, egg cells were carefully opened using a micro spatula. Eggs were counted and summed for each cage.

### Ovarian activation

All workers and queens were dissected under stereomicroscope. The three largest ovarioles were measured using an ocular scale (mm) and at least one ovariole per ovary was measured as in previous studies (14, 23). Mean terminal oocyte length was used in further analyses.

### Fat body lipid analysis

Worker and queen fat bodies were dissected out and homogenized in 5 ml of 2% sodium sulfate, as in (14). 200 μl of the homogenate were transferred to a new vial, with 2.8 ml of a 1:1 chloroform-methanol solution. Vials were centrifuged for 5 minutes at 3000 rpm to separate the lower and the upper phases. The supernatant was transferred to a new vial, where 2 ml of distilled water were added, and samples were centrifuged for 5 minutes at 3000 rpm. The bottom layer (lipids) was separated from the remaining fraction and was placed on a heating plate at 100° C to evaporate the solvent. Lipid amount was determined by using a vanillin-phosphoric reaction (14). A standard curve was created using five different concentrations of vegetable oil in chloroform. Absorbance values (OD 525) were determined using Biotek Synergy LX plate reader and converted into micrograms per bee based on the standard curve equation. The amounts of lipids were normalized for the tissue mass, which was measured on electronic scale prior to analysis, and are presented as percentages.

### Gene expression

The effect of gas treatments on gene expression levels was quantified for six candidate genes and three treatments (full hypercapnia, anoxia and control) in the two experiments of workers and in the virgin and mated queens (see Table S1 for sample size). Analysis included three genes that were regulated by CO_2_ in previous studies (*vitellogenin*, *FOXO*, and *PHGP*) (14, 43), and three hypoxia inducible factors (HIFs) that were regulated by the lack of oxygen in previous studies (44, 45). Vitellogenin (*vit*) codes for the major egg yolk protein in the ovaries (46) and has been shown to be upregulated in CO_2_ treated queens compared to the non-treated controls (23). *FOXO* is a conserved transcription factor that regulates insulin and insulin-like growth factor signaling. It has been shown to be downregulated after CO_2_ treatment. Finally, *PHGP* is a gene coding to an antioxidant enzyme and was previously shown (together with other antioxidant enzymes) to be downregulated in queens in response to hypercapnia (14). HIF-1α (*sima*) and β (*tango*) are two units of a transcription factor involved in the response to low oxygen concentration and *fatiga* is part of a family of prolyl hydroxylases, which plays a vital role in the degradation of HIF-1α. All three genes are expected to be regulated in response to anoxia (45). Design of the forward and reverse primers for each gene was done using primerBLAST. The list of primers and accession numbers is provided in Table S2. Genes with multiple variants had primers designed to target the most conserved region between the sequences. RNA was extracted from queen and worker heads using the RNeasy mini kit according to the manufactural instructions (Qiagen, Valencia, CA). Queens’ RNA was extracted from individual samples whereas heads of workers from the same cage were pooled together. 600 ng of RNA was used for cDNA synthesis with Reverse Transcriptase (Applied Biosystems). After PCR, cDNA was diluted using 85 μl of nuclease free water and stored at −20° C until real time PCR was performed. Gene expression was measured using QuantStudio 5 real-time PCR system (Applied Biosystems). 2μ of the diluted cDNA were used in a reaction with 5 μl SYBR Green, 0.2 μl of the forward and reverse primers, and 4.6 μl of nuclease free water. *Arginine kinase* and *Phospholipase A2* were used as housekeeping genes to control for PCR efficiency and differences across samples. These control genes have been used in previous studies with *Bombus impatiens* (47). Each reaction was performed in triplicate and averaged for data analysis. Each plate had a water control and a negative control (cDNA reaction without Reverse Transcriptase enzyme). Expression levels of each gene were normalized using the geometric means of the two housekeeping genes using the 2^−ΔΔCt^ technique.

### Statistical Analysis

All statistical analyses were performed in JMP Pro 16. The time to enter anesthesia, the recovery time following anesthesia and the number of abdominal contractions were compared using Two-way ANOVA mixed model with caste (virgin, mated, workers) and treatment (FH, PH, and AN) as fixed effect and colony and cage as random factors. The sum of aggressive behavior in workers was compared using Kruskal Wallis test followed by multiple comparison test for each pair using Wilcoxon method. Ovarian activation, fat-body lipid amount, the total egg production and gene expression levels were compared using ANOVA mixed model with colony as a random effect (in queens) or with colony and cage as random effects (in workers). Gene expression data were log transformed prior to analysis. Post-hoc comparison was performed using Tukey HSD test. Statistical significance was accepted at α=0.05. Data are presented as means ± S.E.M

## Results

The time it took bees to become immobile, the recovery time from the gas treatments and the number of abdominal contractions bees exhibited upon recovery were only observed following recovery from anoxia (AN), full and partial hypercapnia (FH, PH), because hypoxia (HY) or control did not result in anesthesia. The time to enter anesthesia differed between all treatments (Two-way ANOVA Mixed model, F_2,158_=134.3, p<0.0001 followed by Tukey HSD p<0.0001) and between workers and virgin queens (F_2,158_=5.18, p=0.006 followed by Tukey HSD p=0.004). Losing mobility was the fastest following FH compared to the other treatments, and workers lost mobility quicker than queens following FH but slower than queens following AN (Fig. 1, Table S3 for means, S.E.M and statistical tests). The recovery time following anesthesia also differed between all treatments (F_2,158_=72.6, p<0.0001 followed by Tukey HSD p<0.0001) and between workers and queens (F_2,158_=50.1, p<0.0001 followed by Tukey HSD p<0.001). Workers were faster to recover compared to queens, but recovery from FH took longer than the other treatments. Finally, we quantified the number of abdominal contractions (AC) in 20 minutes beginning with the first movement across treatments and castes. Here, differences were found between treatments (F_2,158_=14.8, p<0.0001 followed by Tukey HSD p<0.03 for all groups) but not between castes (F_2,158_=1.16, p=0.31), with the highest AC following PH, then FH and the lowest AC following AN.

**Figure 1.**
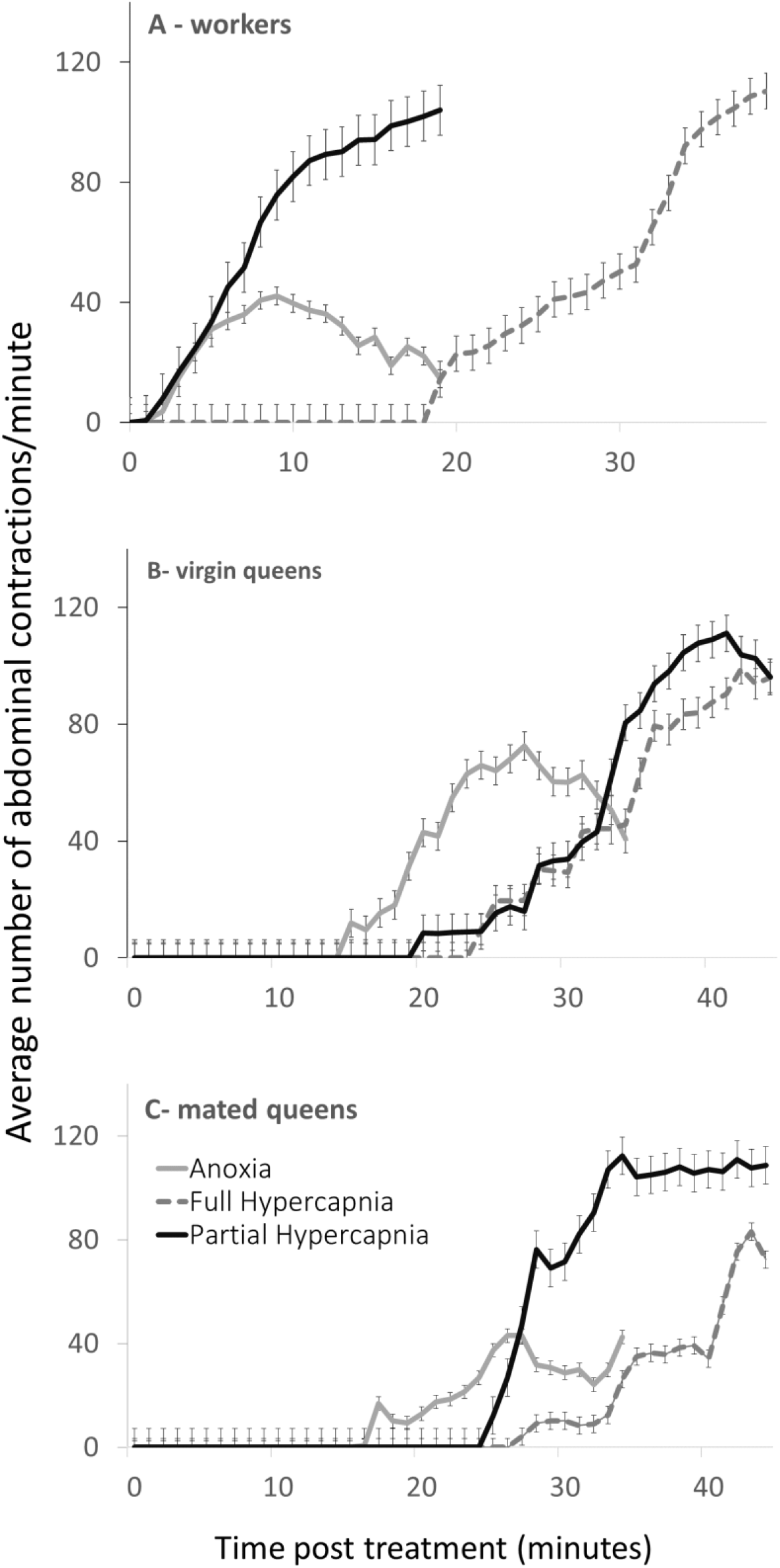
The recovery time from anesthesia and the effect of gas treatments on the number of abdominal contractions in workers (A), virgin queens (B), and mated queens (C). Bees were observed for 20 minutes from their recovery (first movement) in the three treatments that induced anesthesia. Data are presented as means ± S.E.M. Additional data and statistical analyses are provided in Table S3

The effect of the gas treatments on aggression was examined in groups of workers that were treated a single time on the day of emergence. Comparison of all groups showed nearly-significant increase in aggression following CO_2_ (Kruskal Wallis test, *x*^2^_4_=8.98, p=0.06 followed by significant differences between FH and the treatments PH (p=0.037), HY (p=0.004) and control (p=0.029), but not with AN (p=0.07) (Fig. 2).

**Figure 2.**
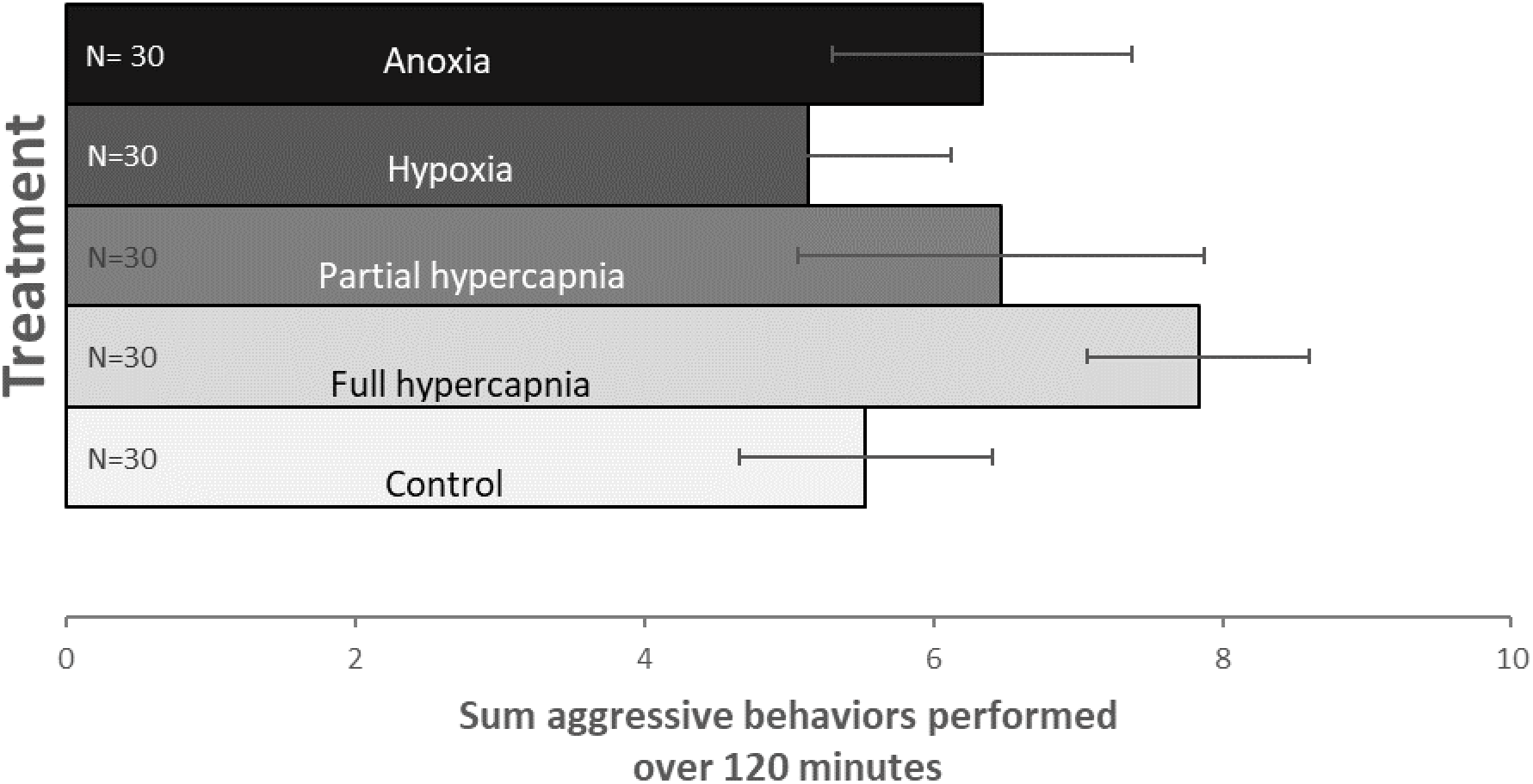
The effect of treatment on the sum aggressive behaviors performed by workers during days 1-6 following emergence. Workers were assigned to five gas treatments on the day they emerged. They were kept in groups of 3 (n=10/treatment) and were observed daily for 20 minutes. Four aggressive behaviors were documented and summed for each worker. Data are presented as means ± S.E.M.

Worker ovary activation was not affected by a single treatment of CO_2_ (ANOVA Mixed model, F_4,140_=0.25, p=0.96, Figure 3A), but was affected by the treatment in the multiply-treated workers (ANOVA Mixed model, F_4, 70_=20.2, p < 0.0001; followed by a Tukey post-hoc test p<0.001 for FH vs. HY and control, PH vs. HY and control, and p<0.04 for An vs. PH, HY and control, Fig 3B). Ovarian activation was also affected by the treatment in the virgin queens (ANOVA Mixed model, F_4,95_=19.3, p<0.001, Fig 3C), where both full and partial hypercapnia treatments differed significantly from the remaining treatments (p<0.001), but not from the anoxia treatment (p>0.05) which displayed intermediate levels of ovarian activation as compared to hypercapnia and control. In mated queen, the data show a similar pattern to the ones of virgin queens with significant effect of the treatment (ANOVA Mixed model, F_4,87_=4.7, p=0.0013 followed by Tukey post-hoc test between=0.03 for AN vs. FH; p=0.009 for FH vs. control, and p<0.001 for FH vs. HY (Fig. 3D). Egg laying was observed in all 10-days old worker cages that were treated once regardless of treatment (n=50), but only in a small fraction of the queen cages (7.7% of all queens, n=154, data not shown). This has no practical meaning since queen-less workers start laying eggs within approximately 7 days from emergence, while queens require 2-3 weeks to initiate a colony following CO_2_ treatment. Thus, 10-days old workers had more time to lay eggs compared to the queens who were sampled 10 days after the gas treatments. Egg laying in workers did not significantly differ across treatments (ANOVA Mixed model, F_4,45_=1.8, p=0.15). In mated queens, two queens laid eggs (in full hypercapnia and anoxia treatments) and in virgin queens, ten queens laid eggs, eight of them in the full and partial hypercapnia treatments. These results are in line with the stronger impact of CO_2_ on the virgin queen ovaries compared to the mated queens.

**Figure 3.**
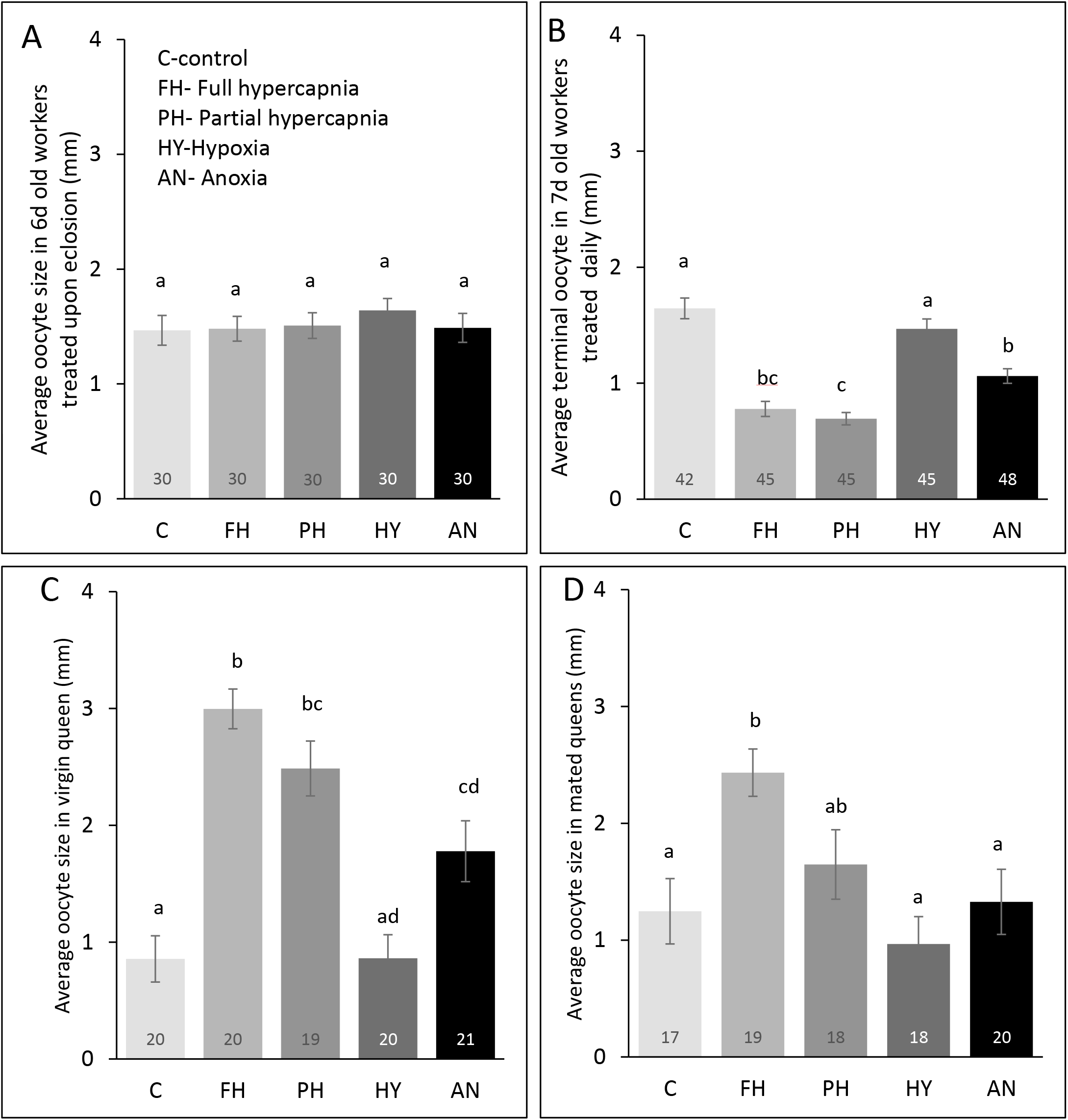
The effect of gas treatment on the average terminal oocyte size in All queens were frozen 10 days post treatment and were 17-20 days old. The sample size per treatment is denoted using the numbers within columns. Different letters denote significant differences at α=0.05. Data are presented as means ± S.E.M.

Lipid percentage in the fat body was overall lower in workers (on average of 4%) compared to queens (on average of 20%). The treatment did not affect the lipid percentage of workers who were treated once (ANOVA Mixed model, F_4,45_=1.3, p=0.297, Fig. 3a), but did affect workers who were treated multiple times (ANOVA Mixed model, F_4. 30_ =3.79, p=0.013 followed by a Tukey post-hoc test p=0.05 between AN and PH vs. C; and p=0.02 between FH vs. C, Fig. 3b). Virgin and mated queens showed similar pattern of response to the treatments with lower lipid percentages in FH and PH compared to the other treatments, but the differences were significant only in the virgin queens (ANOVA Mixed model, F_4,59_=3.8, p=0.009 followed by lower lipids in FH compared to control (p=0.023), AN (p=0.042) and HY (p=0.035), fig. 4C), while the differences in the mated queens were insignificant (ANOVA Mixed model, F_4,45_=1.77, p=0.15, Fig 4D).

**Figure 4.**
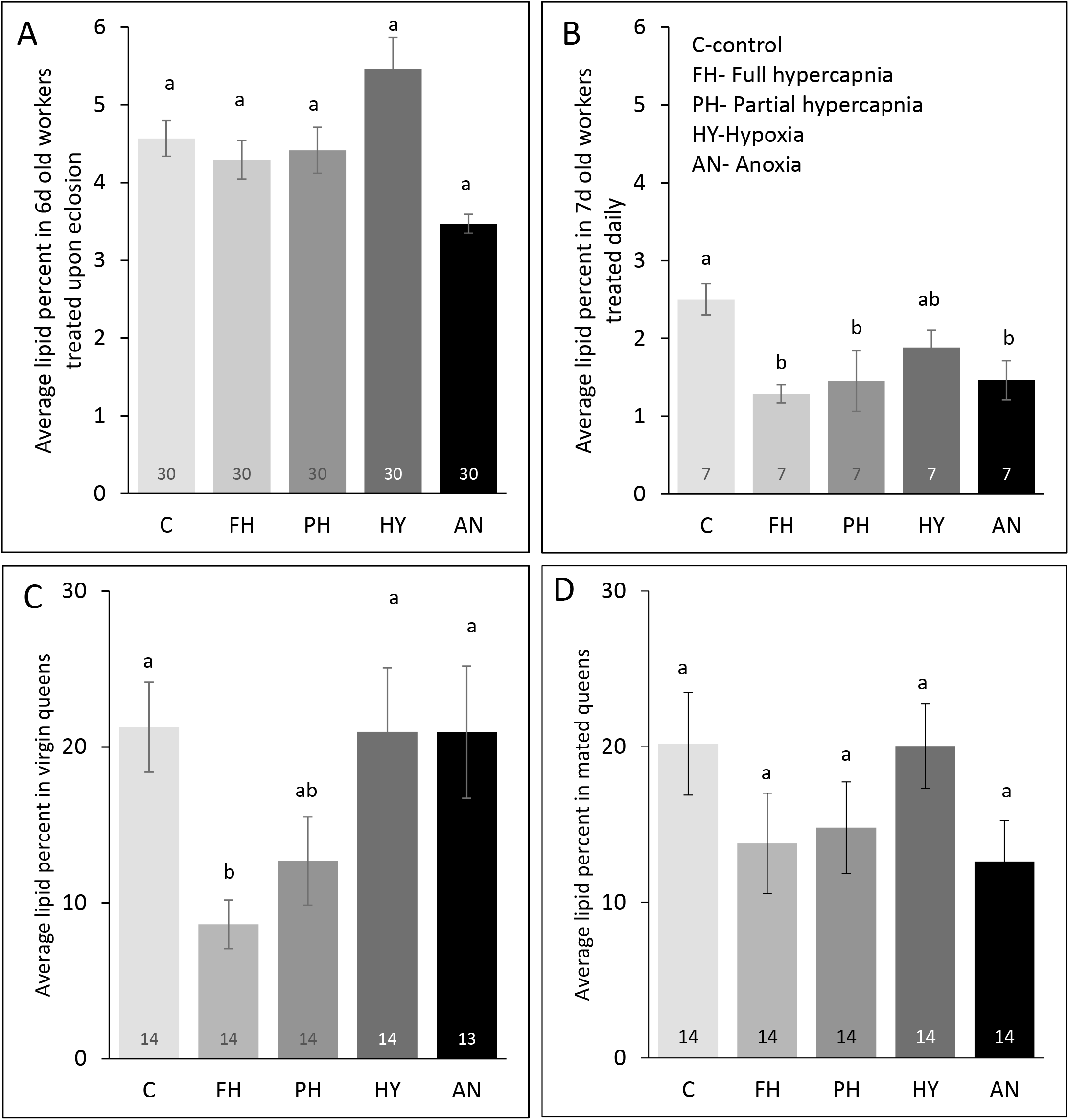
The effect of treatment on the average fat body lipid percent in (A) 6-day-old workers treated upon emergence, and (B) 7-day-old workers treated for six consecutive days after emergence, (C) virgin queens, and (D) mated queens. All queens were frozen 10 days post treatment and were 17-20 days old. The sample size per treatment is denoted using the numbers within columns. Different letters denote significant differences at α=0.05. Data are presented as means ± S.E.M.

Gene expression of six candidate genes, previously shown to be regulated by CO_2_ (*vitellogenin*, *FOXO* and *PHGP*), or by anoxia (*sima, tango, and fatiga*) (45) were tested across treatments in workers, virgin and mated queens. Analysis focused on comparing the control, full hypercapnia, and anoxic treatments. In workers, there was no significant effect of treatment on either the HIF or the CO_2_ genes. This was the case in workers that were treated once (ANOVA Mixed model for *fatiga*: F_2,15_=1.45, p=0.27; for *tango*: F_2,15_=0.55, p=59; for *sima*: F_2,15_=1.16, p=0.34; for *PHPG*: F_2,15_=0.8, p=0.46; for *FOXO*: F_2,15_=0.27, p=77; for *vitellogenin*: F_2,15_=1.1, p=0.35, data not shown), and also in workers that were treated multiple times (ANOVA Mixed model for *fatiga*: F_2,14.4_=1.98, p=0.17; for *tango*: F_2,14.2_=2.6, p=0.11; for *sima*: F_2,14.5_=2.1, p=0.16; for *PHGP*: F_2,12.6_=3.5, p=0.06; for *FOXO*: F_2,14.5_=2.2, p=0.15; for *vitellogenin*: F_2,13.4_=0.41, p=0.67, Fig. 5A). Queen groups were combined, as there was no effect of mating on gene expression levels. In queens, there was a significant effect of treatment on gene expression level in each of the HIF genes (ANOVA Mixed model for *fatiga*: F_2,30_=3.5, p=0.04; for *tango*: F_2,29_=3.5, p=0.04; for *sima*: F_2,27_=9.08, p=0.0009). Additionally, there were significant effects of treatment on the expression levels of *FOXO* and *PHGP*, but not *vitellogenin* (ANOVA Mixed model for *PHGP*: F_2,20_=4.8, p=0.02; for *FOXO*: F_2,29_=5, p=0.01, Fig. 5B).

**Figure 5.**
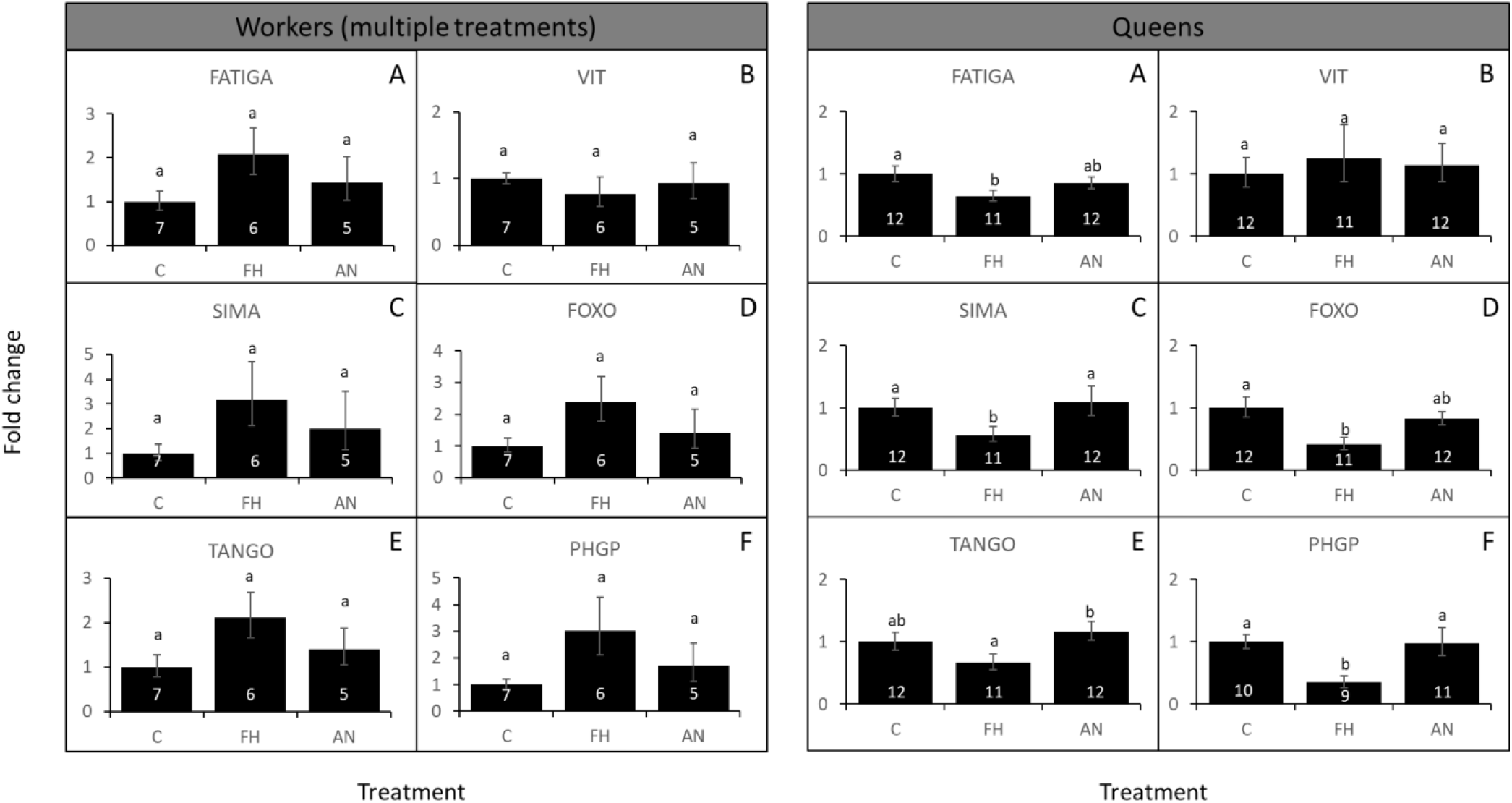
The effect of treatment on the expression level of six genes in *Bombus impatiens* workers (a) and queens (b). Workers were treated daily on days 1-6 after emergence and frozen on day seven. All queens were frozen 10 days post treatment and were 17-20 days old. No differences were found in gene expression of virgin and mated queens and therefore results were combined. The sample size per treatment is denoted using the numbers within columns. Different letters denote significant differences at α=0.05. Data are presented as means ± S.E.M.

## Discussion

In this study, we examined the behavioral, physiological and gene expression effects caused by different gas treatments in *Bombus impatiens* workers and queens. Our goal was to determine if the effects caused by CO_2_ narcosis are mediated through anoxia. Our data demonstrate that overall, this is not the case. Hypercapnia, but not hypoxia or anoxia, caused immobilization in a dose-dependent manner and the recovery from hypercapnia, but not from anoxia, was associated with increased abdominal contraction behavior. Hypercapnia also had marginal impacts on aggression in workers. Physiologically, hypercapnia affected ovarian activation and fat-body lipids much strongly than hypoxia or anoxia. And finally, gene expression differences were significant only in queens, likely due to the smaller sample size in workers. Contrary to our expectations, the three genes associated with anoxia (*fatiga, tango* and *sima*) were downregulated in queens following hypercapnia, but not following anoxia. And, as expected, two out of the three genes associated with CO_2_ in previous studies (*FOXO* and *PHGP*) were significantly downregulated after hypercapnia in queens. Overall, the behavioral, physiological and gene expression differences following hypercapnia were unique to CO_2_ and were not induced by anoxia. The only exceptions were the ovaries of virgin queens that were significantly reduced by both CO_2_ and anoxia, and lipid amount in the multiply-treated workers that were reduced also following anoxia.

A second goal of this study was to understand the impacts of CO_2_ across female castes and following mating. Here we discovered that the recovery from hypercapnia was slower in queens compared to workers, and that also workers exhibit increased aggression following CO_2_ narcosis (14), likely due to their increased ovary activation and competition. We found that full hypercapnia had a stronger effect compared to partial hypercapnia, and that CO_2_ impacts on ovary activation were opposite in queens and workers, and also weaker following mating in queens. We also noted that while queens respond to CO_2_ narcosis, regardless of their age, duration, and number of exposures (14), workers did not respond to CO_2_ upon emergence and require multiple exposures. Overall, the effects of CO_2_ were dose-, caste-, and mating status dependent. These results clarify the diverse effects of CO_2_ in bumble bees, their regulation by caste and mating, and preclude anoxia as a potential mechanism explaining how CO_2_ operates.

Our findings demonstrated that the effects of CO_2_ narcosis are not mediated through anoxia, meaning that CO_2_ has a unique impact on cells, leading to a reduction in extracellular pH either by blocking glutamate receptors in the neuromuscular junction (28, 35) or by altering the levels of other neurotransmitters and/or neuromodulators in the nervous system. Studying these questions requires experimentation at the cell level rather than at the level of the entire organism. Other neurotransmitters and neuromodulators that could be directly affected by CO_2_ are endogenous biogenic amines (BAs) such as dopamine, octopamine, serotonin and tyramine. These low molecular substances are not only involved in reproduction and ovipositing in numerous insects (48–56), but were also shown to be directly affected by CO_2_. In honey bee queens and workers, CO_2_ narcosis was found to affect level of BAs in the brain (36) and also BA receptors in both the ovaries and the brain (57). Selected BAs such as dopamine also correlate with reproductive status in bees in a caste-dependent manner (55). Finally, BA levels can also modify the level of JH (58) and explain its elevated levels following CO_2_ narcosis (23).

Understanding these mechanisms can also clarify why the impacts of CO_2_ are caste dependent in both bumble bees (current study) and honey bees (15, 25, 59). If CO_2_ activates network of genes related to hormones and vitellogenesis that have decoupled during the evolution of social behavior (60–62), then the downstream effects on worker and queen reproduction are easier to explain. For example, if CO_2_ regulates JH and JH decoupled during the evolution of sociality to regulate reproduction in honey bee queens and task allocation in honey bee workers, its caste-dependent effects are intuitive. However, while this explanation makes sense for honey bees, it does not make sense in bumble bees, where JH maintained its gonadotropin role in both queens and workers.

In contrast to its caste-dependent impact on reproduction, CO_2_ causes a reduction in lipids in the fat body of both queens and workers. Typically in insects that reproduce, lipid stores correlate negatively with reproductive status, as lipids are reallocated to build the oocytes (63). However, workers of social species who remain sterile may have low lipid reserves if they use them for maintenance tasks that require energy, like foraging (64–66). Foragers show lower reproductive capacity compared to house bees in both the honey bee (65) and bumble bees (60, 67). This may explain how the same impact of CO_2_ on lipid levels can translate into different reproductive status in queens and workers. It also emphasizes that CO_2_’s primary effect on insect physiology is on metabolism with reproduction being a secondary byproduct of it (Barie, Levin, Amsalem, submitted). The metabolic shift caused by CO_2_ was demonstrated in several species (14, 68, 69). Focusing on the molecular mechanisms that affect metabolism and macronutrient allocation may narrow down the possible mechanisms underlying CO_2_ to explore. One candidate is the insulin signaling pathway that could be triggered by neural changes and affect metabolism.

The weaker effects of CO_2_ on mated queens were observed across all the parameters examined in the study. Post-mating changes occur widely in insects (70–72) and include changes in immunity (71, 73), stimulation of egg laying (74), and a decrease in the response to sex pheromones (70). Since CO_2_ also induces some of these changes in bumble bees (14), its impact following mating may be redundant. It should be noted that the only effect of anoxia in this study was observed in the ovary of virgin queens which could reflect a higher sensitivity to lack of oxygen during earlier life stages. Interestingly, response to anoxia is indeed graded, tissue and age dependent (75).

Overall, our data show that the behavioral, physiological and gene expression differences following hypercapnia were unique to CO_2_ and were not induced by anoxia, with the exception of ovarian activation being affected by both hypercapnia and anoxia in virgin queens. We also demonstrated that CO_2_ affects queens and workers differently alongside a conserved effect of CO_2_ on lipid level in both castes and likely also across species. Our results contribute to understanding of the mechanisms of CO_2_ narcosis in bees. Further studies at the cell level could help identify the molecular mechanisms underpinnings CO_2_ effect on the neural system and how these are translated into metabolic differences in insects.

## Acknowledgements

We thank members of the Amsalem Lab for the thoughtful discussion and critical reading of earlier drafts of the manuscript. We also thank undergraduate assistant Angelina Toral for her assistance with this experiment.

## Author’s contributions

AC and EA designed the experiments, AC collected and analyzed the data, AC and EA wrote the manuscript.

