## Supplemental Tables S1-S3 for "Impacts and mechanisms of CO_2_ narcosis in bumble bees: Narcosis depends on dose, caste and mating status and is not induced by anoxia"

### Supplementary materials

**Table S1**: The sample size of bees in the study split into the different treatments and variables

| | | Abdominal contractions (bees) | Aggression (bees) | Ovary activation (bees) | Egg laying (cages) | Fat-body lipids (bees) | Gene expression (bees) |
|---|---|---|---|---|---|---|---|
| Worker – single treatment (6d) | C | X | 30 | 30 | X | 30 | 6 |
| | FH | 39 | 30 | 30 | X | 30 | 6 |
| | PH | 30 | 30 | 30 | X | 30 | 6 |
| | HY | X | 30 | 30 | X | 30 | 6 |
| | AN | 30 | 30 | 30 | X | 30 | 6 |
| Worker – single treatment (10d) | C | X | X | X | 10 | X | X |
| | FH | X | X | X | 10 | X | X |
| | PH | X | X | X | 10 | X | X |
| | HY | X | X | X | 10 | X | X |
| | AN | X | X | X | 10 | X | X |
| Workers – multiple treatments (7d) | C | X | X | 42 | X | 7 | 7 |
| | FH | X | X | 45 | X | 7 | 6 |
| | PH | X | X | 45 | X | 7 | X |
| | HY | X | X | 45 | X | 7 | X |
| | AN | X | X | 48 | X | 7 | 5 |
| Virgin queens | C | X | X | 20 | 20 | 14 | 6/4* |
| | FH | 10 | X | 20 | 20 | 14 | 5/3* |
| | PH | 12 | X | 19 | 20 | 14 | X |
| | HY | X | X | 20 | 20 | 14 | X |
| | AN | 11 | X | 21 | 21 | 13 | 6/5* |
| Mated queens | C | X | X | 17 | 10 | 14 | 6/6* |
| | FH | 10 | X | 19 | 11 | 14 | 6/6* |
| | PH | 10 | X | 18 | 11 | 14 | X |
| | HY | X | X | 18 | 10 | 14 | X |
| | AN | 8 | X | 20 | 11 | 14 | 6/6* |

*The first number applies to all genes whereas the second number applies to the gene PHGP, where four samples omitted due to a technical problem.

**Table S2**: primers sequences and accession numbers of *Bombus impatiens* used in the study.

| Gene | Accession no. | F primer | R primers |
|---|---|---|---|
| *Phospholipase A2 (HKG)* | LOC100747217 | CATTTCCGCAAGTGGTAGGT | GGTCACACCGAAACCAGATT |
| *Arginine kinase (HKG)* | LOC100741837 | GTTGGTAGGGCAGAAGGTCA | AGGTCTACCGTCGTCTGGTG |
| *sima* | LOC100746032 | GGAAAAGTCTCGTGATGCTGC | AGATGCCTGTTCCGATGGC |
| *tango* | LOC100740860 | GAAGCACCTGATTCTGGAGGC | TGTAATTCAAAACAGGCGCCAC |
| *fatiga* | LOC100747684 | CGGACCGATAGTCAGTCGTG | GCACCGTGAACACCTTAGGA |
| *Vitellogenin* | LOC10074717 | CAGCCGCCAATATGATACCT | CCCTCCGTTCGAAGTGATAA |
| *FOXO* | LOC100745007 | CAACAACAATCGCAAACAGG | CTCCCATCAATTGTCCCATC |
| *PHGP* | LOC100745584 | ACTGTACGATGAGTATGCTGAC | TTCGGTACCTCCTGGTTCCT |

**Table S3**: The time to enter anesthesia, the recovery time from anesthesia and the number of abdominal contractions exhibited by bees in 20 minutes from the initiation of movement. Data are presented as means ± S.E.M

| | Workers | Virgin queens | Mated queens |
|---|---|---|---|
| Time to enter anesthesia (seconds) | | | |
| Full hypercapnia | 14.3±0.3 | 17.8±0.7 | 18.7±0.4 |
| Partial hypercapnia | 34.7±1 | 29.8±2.2 | 31.2±1 |
| Anoxia | 29.9±0.9 | 21.5±1.2 | 22.3±1.3 |
| Effect of treatment | ANOVA Mixed model, $F_{2,158}$=134.3, p<0.0001 followed by Tukey HSD p<0.0001 for all groups | | |
| Effect of caste | ANOVA Mixed model, $F_{2,158}$=5.18, p=0.006 followed by Tukey HSD p=0.004 for workers vs virgin queens | | |
| The recovery time from anesthesia (minutes) | | | |
| Full hypercapnia | 29.7±1 | 33.9±1.9 | 38.4±0.7 |
| Partial hypercapnia | 7.07±0.6 | 31.5±1.4 | 28.8±0.8 |
| Anoxia | 4.33±0.4 | 21.6±1.1 | 24.6±2.2 |
| Effect of treatment | ANOVA Mixed model, $F_{2,158}$=72.6, p<0.0001 followed by Tukey HSD p<0.0001 for all groups | | |
| Effect of caste | ANOVA Mixed model, $F_{2,158}$=50.1, p<0.0001 followed by Tukey HSD p<0.001 for workers vs. queens | | |
| Abdominal contractions number in 20 minutes | | | |
| Full hypercapnia | 1215±116 | 1190±200 | 594±161 |
| Partial hypercapnia | 1264±162 | 1384±183 | 1764±73 |
| Anoxia | 526±42 | 955±198 | 475±170 |
| Effect of treatment | ANOVA Mixed model, $F_{2,158}$=14.8, p<0.0001 followed by Tukey HSD p<0.03 for all groups | | |
| Effect of caste | ANOVA Mixed model, $F_{2,158}$=1.16, p=0.31 | | |
